# Proviral dynamics and HIV-1C viral diversity in the context of HIV-TB co-infection

**DOI:** 10.64898/2026.04.08.712756

**Authors:** Shilpa Bhowmick, Sharad Bhagat, Sapna Yadav, Kartik Kadam, Prajakta Kamble, Shweta Shrivas, Pratik Devadiga, Snehal Kaginkar, Varsha Padwal, Namrata Neman, Satyajit Musale, Nandan Mohite, Vidya Nagar, Priya Patil, Sachee Agrawal, Sushma Gaikwad, Jayanthi Shastri, Nupur Mukherjee, Kiran Munne, Vikrant M. Bhor, Taruna Madan, Jyoti Sutar, Jayanta Bhattacharya, Vainav Patel

## Abstract

**Background:** ART effectively suppresses HIV replication and restores CD4+ T cells; however, long-lived HIV latent reservoirs enable viral persistence. Tuberculosis (TB) co-infection further impacts HIV latency and enhances viral replication. Given the high prevalence of latent TB infection (LTBI) in TB-endemic settings, understanding its impact on HIV biology is critical. Our study aims to investigate the influence of TB co-infection on HIV reservoir dynamics, viral diversity, and drug resistance mutations in ART-naive individuals.

**Methodology:** Samples from 90 ART-naive HIV-1C individuals, stratified based on IGRA and TB diagnosis, were used in this study. Plasma and PBMCs were isolated for viral RNA and DNA extraction respectively. Total proviral DNA was quantified using *gag* PCR. Full-length *env* and *pol* genes were amplified, purified and sequenced using ONT and Illumina platforms. *Pol* sequences were subjected to Drug Resistance Mutation (DRM) analysis via Stanford HIVdb with a minimum threshold mutation frequency of ≥10%. Full length *env* sequences were used for phylogenetic analysis by aligning with Indian Subtype C reference sequence and phylogenetic tree was generated using ggplot2.

**Result:** Proviral load analysis showed no significant differences across HIV+LTBI−, HIV+LTBI+, and HIV+TB+ groups, although a trend toward higher levels was observed in HIV+TB+ individuals. Correlation analysis revealed distinct immune associations, with HIV+LTBI+ individuals showing positive correlations with activation and PD-1 expression. Longitudinal analysis of proviral loads demonstrated a modest decline in proviral load post-ART but remained persistent for up to 18-20 months following initiation of ART accompanied by low level ongoing viral replication. DRM analysis revealed a 33% prevalence in ART-naive individuals, with higher occurrence in HIV+LTBI+ group. Of the identified DRMs, 38% (5/13) and 71% (5/7) in sequences obtained from PBMC and plasma respectively were attributed to polymorphic mutations associated with Integrase strand transfer inhibitors (INSTIs). DRMs within plasma and PBMC derived viruses showed high concordance. Phylogenetic analysis of *env* sequences indicated overlapping viral populations between the 3 groups, with greater diversity in PBMCs compared to plasma.

**Conclusion:** The study highlights that HIV reservoir dynamics, drug resistance, and viral diversity are significantly influenced by TB co-infection. While proviral loads were comparable, LTBI-associated immune activation and granuloma niches may have driven viral diversification and DRM emergence. High concordance between compartments and presence of transmitted resistance underscore the need for baseline screening, multi-compartment analysis, and sustained surveillance.

## Introduction

Antiretroviral therapy (ART) has transformed HIV into a manageable condition by suppressing plasma viremia, limiting viral sequelae, and enabling recovery of CD4⁺ T cells through inhibition of key viral enzymes (1). Despite this, HIV persists due to the establishment of long-lived latent reservoirs composed of replication-competent proviruses integrated into the host genome, which remain transcriptionally silent and are not eliminated by anti-retroviral therapy (ART) (2–4). Tuberculosis co-infection further perturbs this equilibrium by disrupting latency and promoting viral reactivation. Mechanistically, Mycobacterium tuberculosis components, such as phosphatidylinositol mannoside 6, activate pattern recognition receptors including TLR-2, leading to MyD88-dependent NF-κB signalling that enhances HIV transcription (5). Consistently, both experimental and clinical studies demonstrate that HIV–TB co-infected individuals exhibit higher viremia and increased cell-associated HIV-1 DNA compared to HIV mono-infected individuals, indicating that TB can destabilize viral reservoirs and accelerate disease progression (2,6).

Additionally, HIV-1 viral diversity continues to present significant obstacles for both prevention and treatment predominantly due to its remarkable genetic heterogeneity. This heterogeneity is a consequence of the virus’s inherently error-prone reverse transcriptase, its rapid replication rates, the phenomenon of recombination, and the persistent immune selection occurring within the host organism (7,8). Concurrently, co-infections play an instrumental role in modulating HIV biology. Tuberculosis (TB) stands as the highest comorbidity among PLHIV (9). Evidence shows that TB does not merely coexist with HIV but actively facilitates viral replication and amplifies viral heterogeneity (10). Investigations report that active HIV–TB co-infection is associated with heightened immune activation and T cell exhaustion compared to HIV mono-infection, driven by elevated inflammatory mediators such as IP-10, D-dimer, TNF-α, and MCP-1 (11,12), which might further enhance HIV replication. Also, HIV-1 spreads to various anatomical sites where the site-specific microenvironment regulates its evolutionary kinetics (13). Latent TB infection, affecting 20-60% of Indian population (14,15), consists of granulomatous lesions harboring cells co-infected with MTB and HIV might present as a niche for viral evolution within these individuals (16–18). Collectively, these findings reveal a complex connection wherein tuberculosis-induced immune activation influence viral replication, quasi-species diversity, and thus vulnerability to anti-retroviral therapy and bnAb-mediated neutralization. Understanding this intricate relationship and targeting the HIV latent reservoir is vital for the formulation of efficacious antibody-based prevention and therapeutic strategies in contexts of HIV-TB coinfections. In this study we undertook analysis of reservoir dynamics, associated DRMs and viral diversity in ART naïve HIV-1C infected individuals representing different states of TB co-infection.

## Materials and methods

### Ethics statement

The study received approval from the ICMR-NIRRCH Ethics Committee (Ethics No. 348/2018, 410/2020), the ART Centre of the JJ Group of Hospitals (No.: IEC/Pharm/RP/183/Oct/2020), ECARP at B.Y.L. Nair Hospital (ECARP/2020/152 and ECARP/2020/109), as well as NACO. Participants for the study were systematically recruited from JJ Hospital and B.Y.L. Nair Charitable Hospital, Mumbai, in full compliance with the sanctioned protocol. Signed informed consent forms were meticulously acquired from the study participants prior to the commencement of sample collection.

### Study Participants

Ninety ART naïve HIV-1C infected individuals were enrolled in the study, with peripheral blood samples procured in EDTA and lithium heparin vacutainers subsequent to the acquisition of signed informed consent from all participants. HIV Tridot assay was performed to affirm the sero-positivity for HIV-1. Blood samples obtained in lithium heparin tubes were used to perform Interferon Gamma Release assay (IGRA) employing the QuantiFERON-TB Gold Plus at NIRRCH and participants were further stratified into distinct groups as LTBI+ and LTBI-based on IGRA results. The diagnosis of active TB was established through hospital reports via chest radiography, sputum and/or smear positivity, and GeneXpert positivity. HIV-1 positive individuals concurrently infected with LTBI were monitored for duration of up to 18-20 months at six-month intervals following the commencement of anti-retroviral therapy (ART). Time point zero (TP0) signifies the initial cross-sectional sampling. Time point one (TP1) denotes the blood collection following 6-8 months of ART, time point two (TP2) signifies 12-14 months of ART, and time point three (TP3) corresponds to 18-20 months of ART. Whole blood was acquired at each time point to similar assays.

### Plasma separation and PBMC isolation

Whole blood samples were gently mixed to ensure homogeneity. The blood was then centrifuged at 1200 g for 10 minutes. Following centrifugation, the upper plasma layer was carefully transferred to a fresh 15 mL conical centrifuge tube and centrifuged again at 1200 g for 10 minutes at to remove residual cellular debris. The clarified supernatant was collected and aliquoted into sterile 1.5 mL microcentrifuge tubes as 600 μL plasma fractions. An aliquot of 50 μL plasma was used for the HIV TRI-DOT test and the remaining plasma was snap frozen and stored at −80°C for subsequent isolation of viral RNA.

Peripheral blood mono-nuclear cells (PBMCs) was isolated by density gradient centrifugation method using HiSep LSM 1077. After separation of plasma, the blood was gently mixed by inversion and diluted in a 1:2 ratio with Roswell Park Memorial Institute (RPMI) 1640 medium. Three mL of HiSep LSM 1077 was aliquoted into a sterile 15 mL conical centrifuge tube. The diluted blood sample was carefully layered over the density gradient medium at a ratio of 1:3 (one part of LSM to three parts of diluted blood). Samples were centrifuged at 600 g for 20 minutes at room temperature without brake with minimum acceleration and deceleration. The buffy coat at the interface between plasma and LSM was carefully aspirated using a Pasteur pipette and transferred to a fresh sterile 15 mL conical tube. The volume was adjusted to 10 mL with Dulbecco phosphate buffered saline (DPBS) with 2% FBS and centrifuged at 350 g for 10 minutes at room temperature. The supernatant was discarded to remove residual LSM, and the cell pellet was gently resuspended in 200 μL Dulbecco phosphate buffered saline. An aliquot of 20 μL cell suspension was reserved for DNA extraction.

### Proviral load estimation

The quantification of proviral HIV-1 DNA was performed using HIV-1 specific *gag* PCR as described previously (3). Briefly, 100 ng of DNA template, which corresponds to 15,151 cells (standardized for *gapdh* copies utilizing qPCR) (19), was derived from the PBMC and subsequently subjected to nested *gag* PCR amplification.

Standards for copy number quantification were established by combining 1:10 serially diluted J-Lat 8.4 cells DNA with DNA obtained from uninfected PBMC, thereby maintaining a final DNA concentration of 100 ng. The HIV DNA copies were quantified and reported as copies per million cells.

### Viral and genomic DNA isolation and cDNA synthesis

QIAamp® Viral RNA Mini kit (Qiagen, Cat. no. 52904) was used to isolate viral RNA from plasma samples as per manufacturer’s instruction. Genomic DNA was isolated from peripheral blood mononuclear cells (PBMC) using QIAmp blood DNA mini kit (Qiagen, Cat. no. 51106) as per the manufacturer’s instructions. Plasma isolated RNA was primed with EnvR1 oligo (5′-GCACTCAAGGCAAGCTTTATTGAGGCT-3′) proximal to 3’ end of the HIV RNA genome (HXB2: 9605–9632) and Aenvseq4 (5’-CAAGCTTGTGTAATGGCTGAGG-3’) binding downstream of the pol gene (HXB2: 6817–6838). Synthesis of cDNA was performed using the Superscript III first strand synthesis kit (Invitrogen, Catalog no: 18080051) following the protocol provided by the manufacturer.

### Amplification of full length *gp160* and *pol*

Full-length *env (gp160)* and *pol* genes were amplified using nested PCR approach from HIV-1 infected plasma and PBMC samples as described previously (35). *Rev-env gp160* cassette was amplified from the cDNA and DNA product using La Taq high fidelity DNA polymerase in the 1st round (Takara Bio Inc., Cat no. RR002M) and PrimeSTAR GXL high fidelity DNA polymerase (Takara Bio Inc. Cat no. R050A) in the second round. The primers used for the 1st round was EnvF1: 5′-AGARGAYAGATGGAACAAGCCCCAG-3′ (HXB2: 5550–5574) and EnvRP2: 5′-GTGTGTAGTTCTGCCAATCAGGGAA-3′ (HXB2: 9157–9181) while for the second round were Env IF: 5′-CACCGGCTTAGGCATCTCCTATGGCAGGAAGAA-3′ (HXB2: 5950–5982) and EnvIR: 5′-TATCGGTACCAGTCTTGAGACGCTGCTCCTACTC-3′ (HXB2: 8882–8915). PCR conditions followed for 1st round was initial denaturation of 94°C for 2 min followed by 10 cycles of 94°C for 10 s, 60°C for 30 s, 68°C for 3 min, 15 cycles of 94°C for 10 s, 55°C for 30 s, and 68°C for 3 min with final extension of 68°C for 10 min. PCR conditions followed for 2^nd^ round were initial denaturation of 94°C for 2 min followed by 10 cycles of 94°C for 10 s, 62°C for 30 s, 68°C for 3 min, 20 cycles of 94°C for 10 s, 60°C for 30 s, and 68°C for 3 min with final extension of 68°C for 10 min. The primers used for the 1st round toward *pol* amplification were Pro5F: 5′-AGAAATTGCAGGGCCCCTAGGAA-3′ (HXB2: 1996–2018) and PolR1: 5′-GGTACCCCATAATAGACTGTRACCCACAA-3′ (HXB2: 6324–6352) while for the second round were Pro3F: 5′-AGANCAGAGCCAACAGCCCCACCA-3′ (HXB2: 2143–2166) and PolR2: 5′-CTCTCATTGCCACTGTCTTCTGCTC-3′ (HXB2: 6207–6231). PCR conditions followed in both rounds were initial denaturation of 94°C for 2 min followed by 15 cycles of 94°C for 10 s, 65°C for 30 s, 68°C for 3 min, 20 cycles of 94°C for 10 s, 55°C for 30 s, and 68°C for 3 min with final extension of 68°C for 10 min.

### Purification of full length *gp160* and *pol* amplicons

The second-round PCR products were resolved on a 1% agarose gel, and the band of interest was excised using a sterile scalpel. The excised gel fragment was purified using the NucleoSpin® Gel and PCR Clean-up kit (Macherey-Nagel, Cat. No.740609.50) according to the manufacturer’s instructions.

### Long and short read sequencing of *gp160* and *pol* amplicons

The *env* amplicons were purified and subsequently subjected to long read (Oxford nanopore) deep sequencing to obtain dominant sequences whereas the *pol* amplicons were purified and subsequently subjected to library preparation for short read Illumina sequencing. Libraries for short read Illumina sequencing was prepared using NEBNext Ultra II FS DNA Library Prep Kit for Illumina (Cat no. E7805L) and NEBNext Multiplex Oligos for Illumina (96 Unique Dual Index Primer Pairs) (Cat no. E6440S) and libraries were run using Nextseq 2000 platform Libraries for ONT sequence was prepared using Native Barcoding Kit 96 V14 (Cat no. SQK-NBD114.96) and library was run using Flow-cell R10.4.1 (FLO-MIN114) on MinION Mk1B.

### Anti-retroviral drug resistance mutation (DRM) analysis of *pol* gene

The reads generated from Illumina sequencing of *pol* gene were trimmed using Trimmomatic and checked for their Phred quality. DRM data was generated from sequences having read depth >50 and mutation detection threshold >10% with Contig sequences obtained through residue-wise amino acid mutations (*pol*) using Stanford HIVdb DRM prediction system. DRM datasets thus generated were used to generate Circos plot for presence and frequency of DRM using R studio v2026.01.0.

### Generation of *env* consensus sequence and phylogenetic analysis

Full-length *env* amplicons, 3′ fragments, were sequenced using long-read Oxford Nanopore Technology (ONT) platform. The raw data obtained from the nanopore sequencing were converted to Fastq files using Dorado basecaller (v.1.3.0) followed by demultiplexing. Raw reads were filtered for quality and read length using seqtk, with reads shorter than 300 bp removed to eliminate formatting inconsistencies and retain high-quality long reads for downstream analysis. The reads were then aligned to the HIV-1 subtype C reference sequence (GenBank ID: AF067155.1) using Minimap2 and processed for read sorting and filtration with samtools. Reads were merged together into consensus sequences using using Medaka (v1.x) with the model r1041_e82_400bps_sup_v5.2.0. The consensus sequences obtained were further used for generation of phylogenetic tree. Phylogenetic tree was generated for HIV-1C envelope sequences as described earlier (20). Briefly, the sequence data sets were aligned using MAFFT (v7), and sequences were combined into a single Fasta file. The resulting alignment was trimmed using trimAl (automated1 mode) to remove poorly aligned regions and gaps. The phylogenetic analysis was performed using IQ-TREE (v2) and visualised in R studio using the ggplot 2.

### Statistical Analysis

GraphPad Prism software version 10 was used to perform statistical analysis. The data were represented as violin plots indicated with median values. Comparison between groups was performed by the Kruskal–Wallis one-way ANOVA non-parametric test. Comparisons between PBMC and plasma compartment for matched samples was done using Mann-Whitney t-test for unpaired analysis and Wilcoxon signed rank test for paired analysis. For all statistical calculations, p < 0.05 was considered significant. R studio was used to generate circos plot and phylogenetic tree.

## Results

### Immunological and virological characteristics of Study population

Samples from a total of 90 ART naïve HIV-1C infected individuals, stratified as LTBI negative (HLTBI-[n=46]) or positive (HLTBI+ [n=25]) and active TB positive (HTB+ [n=19]). As described in Table 1, all 3 groups had similar median age, of which the HIV+TB+ group had significantly lower CD4+ T cell counts (HIV+LTBI-, p= 0.081; HIV+LTBI+, p= 0.0001). HIV+LTBI+ group had significantly higher CD4:CD8 ratio compared to HIV+TB+ group (p=0.0007). Conversely, the HTB+ group exhibited the highest levels of viremia, indicating that active TB co-infection may have enhanced HIV-1 replication and was associated with greater disease severity, as reflected by reduced CD4 counts and lower CD4:CD8 ratios.

**Table 1:** Abridged table for HIV disease progression markers at cross-sectional time point.

|  | HIV+LTBI- | HIV+LTBI+ | HIV+TB+ |
| --- | --- | --- | --- |
| No. of participants | n= 46 | n= 25 | n= 19 |
| Gender (F/M) | 16/30 | 7/18 | 5/14 |
| Age (years)<sup>a</sup> | 37(19-58) | 41 (26-59) | 41 (29-59) |
| CD4+ T cell (cells/μL)<sup>a</sup> | 324<br>(6-835) | 381<br>(10-849) | 127*<br>(28-455) |
| CD4/CD8<sup>a</sup> | 0.22#<br>(0.01-1.26) | 0.39<br>(0.02-1.56) | 0.12<br>(0.05-0.5) |
| HIV-1 Viral load (copies/mL)<sup>a</sup> | 40,823<br>(24.91-59,20,711) | 90,982<br>(654-6,21,950) | 2,57,156<br>(66.7-1,11,106,33) |
| HIV-1 Proviral load (copies/million genomic equivalents)<sup>a</sup> | 4,492<br>(183-44071) | 4,298<br>(156-78,630) | 5,973<br>(1086-101183) |

Footnote: <sup>a</sup>Data represented as median (range)
\*CD4+ T cell absolute counts in HIV+TB+ group was significantly lower than other 2 HIV+ groups.

### Proviral load analysis of ART naïve HIV-1 infected individuals at different stages of HIV-TB co-infection

To investigate whether HIV reservoir size varies across different stages of HIV–TB co-infection, proviral DNA levels were quantified in ART-naïve HIV-1 infected individual belonging to three clinical groups. Comparative analysis revealed that there was no statistically significant difference in proviral loads among the three HIV-positive groups (Figure 1A). This observation was further supported by hierarchical clustering analysis (Figure 1B), which did not show a clear separation of individuals based on proviral load across the groups. Albeit, a trend toward higher proviral loads was observed in the HIV+ TB+ group reflecting the heightened immune activation observed during active co-infection, with enhanced viral replication and more advanced disease progression (12).

**Figure 1:**
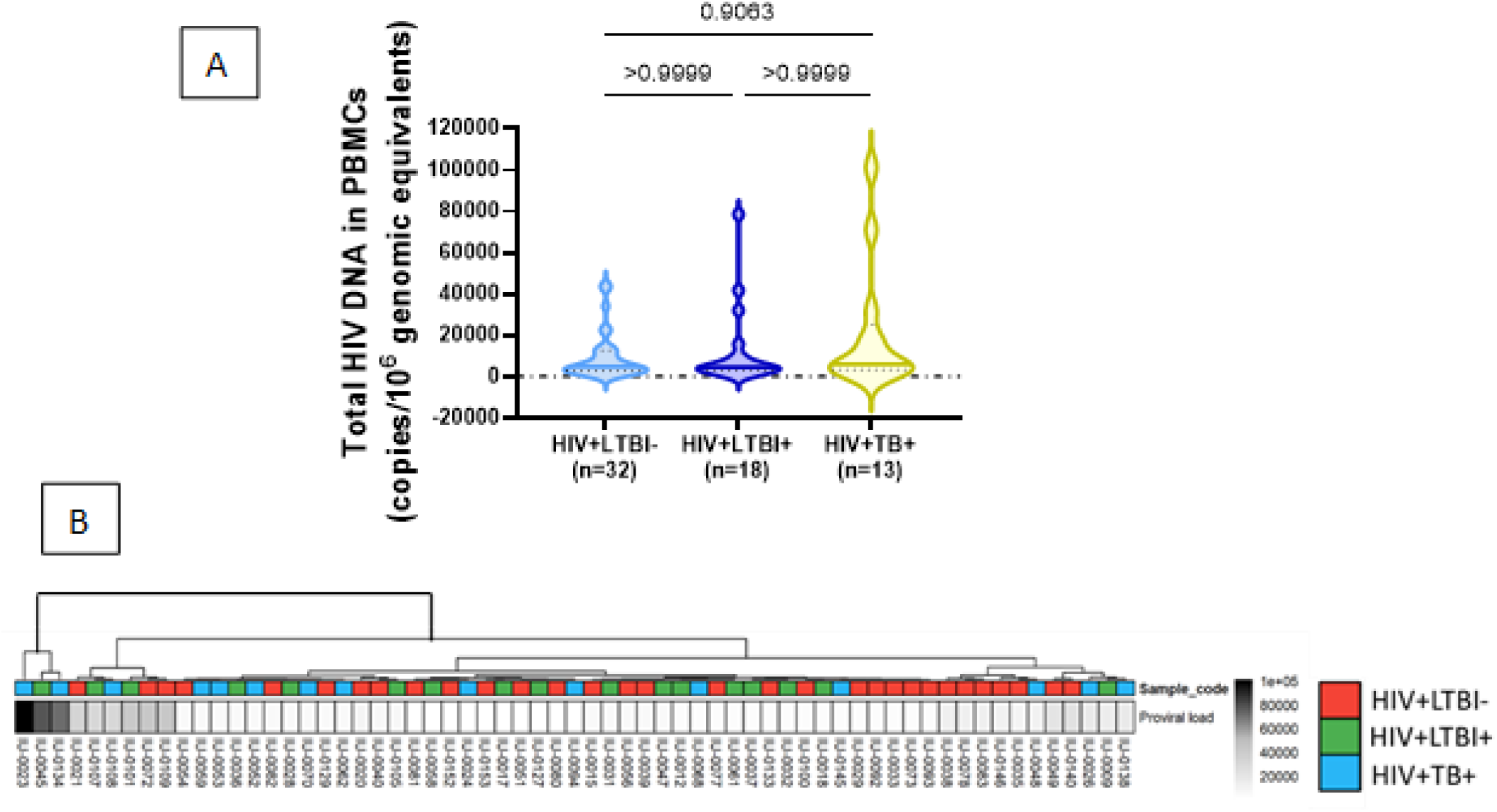
Levels of Proviral load at cross-sectional time point: (A) Difference in proviral loads between HIV+LTBI-(n=32), HIV+LTBI+(n=18) and HIV+TB+(n=13) groups. (B) Hierarchical clustering analysis of proviral loads in ART naïve HIV-1 infected individuals. Comparisons between groups were calculated by Kruskal-Wallis one-way ANOVA non-parametric test.

### Correlational analysis between proviral loads and systemic immune markers in ART naïve HIV-1 infected individuals

To further understand the relationship between host immune responses and HIV reservoir burden, we performed correlation analysis between systemic immune parameters (systemic immune data obtained from *Bhowmick et al, Cells, 2025* (12) study performed on the same cohort) and proviral loads across the 3 study groups (Figure 2). In the HIV+TB+ group, a negative association was observed between regulatory T cells (Tregs) and proviral loads, which might be reflective of markedly reduced CD4⁺ T cell counts, increased immune activation and viral replication in these individuals that could have led to enhanced infection and subsequent death of CD4+ T cells including reservoir-containing cells. A similar negative trend was also observed in the HIV+ LTBI-individuals, although the strength of the association was weaker. Interestingly, in HIV+ LTBI+ group we observed positive association between proviral loads with activation and PD-1 expression indicating a more functional and immunocompetent CD4⁺ T-cell compartment.

**Figure 2:**
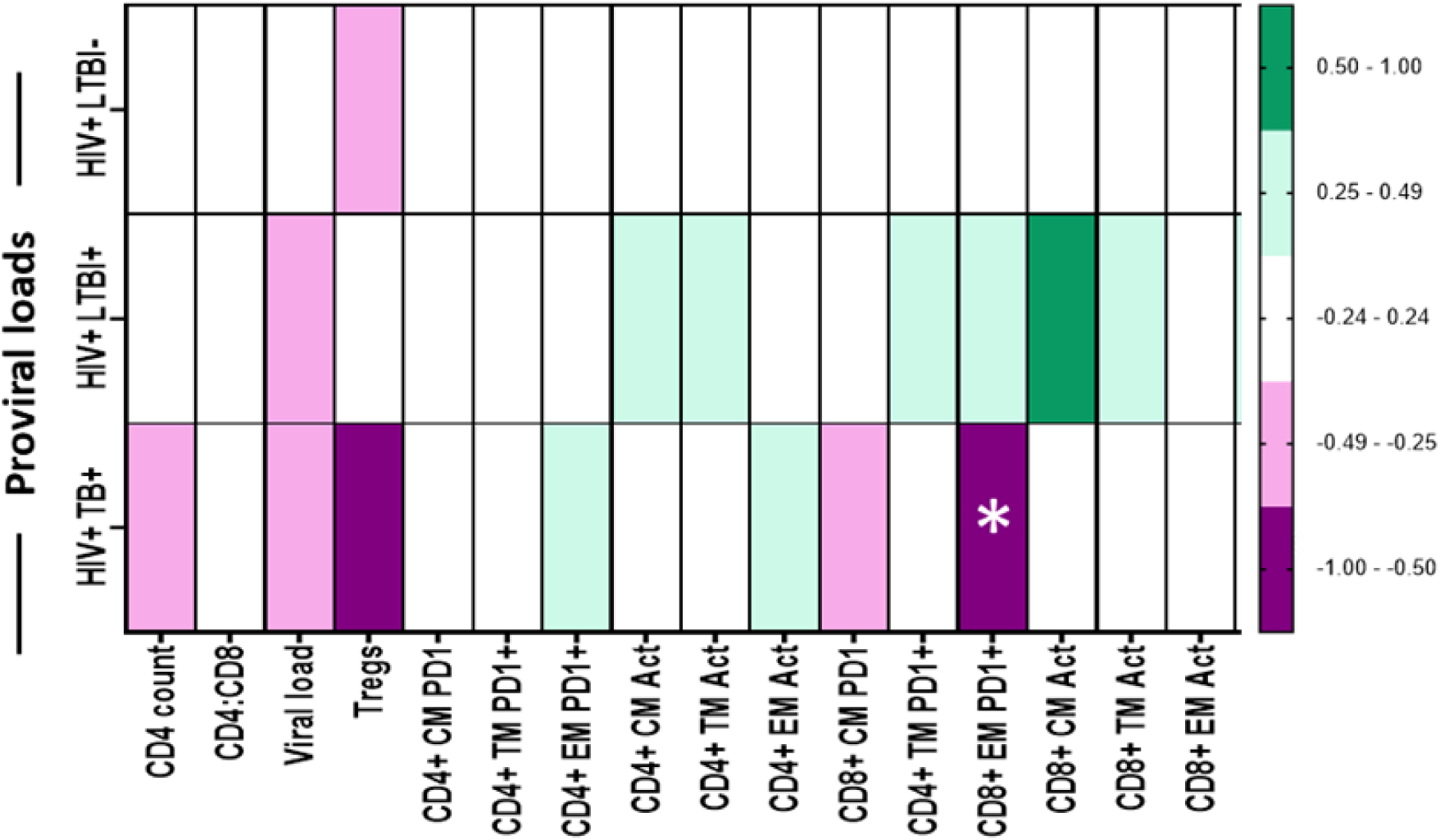
Association between systemic immune responses and proviral loads: Heatmap for correlation between systemic immune response and proviral loads in HIV+LTBI-(n=30), HIV+LTBI+ (n=17) and HIV+TB+ (n=15) groups. p and r values for associations were determined by Spearman’s correlation test.

### Dynamics of Proviral Load Following Initiation of Antiretroviral Therapy in HIV+LTBI+ individuals

To evaluate the dynamics of the HIV-1 reservoir following treatment initiation, proviral loads were assessed longitudinally for up to 18 months in HIV+LTBI+ individuals after the commencement of antiretroviral therapy (ART). A decline in proviral load was observed at the TP1 i.e. 6-8 months following ART initiation (Figure 3 A), indicating a reduction in the reservoir burden during the early phase of therapy. This trend was consistent with the viral load measurements, which also demonstrated a reduction during this period (Figure 3C). However, by TP2 i.e. 12-14 months, the levels of proviral load showed a subsequent increase compared to 6-8 months. Furthermore, analysis of matched longitudinal samples from the same individuals demonstrated a similar pattern, with an initial decline in proviral loads followed by a modest increase at later time points (Figure 3B)

**Figure 3:**
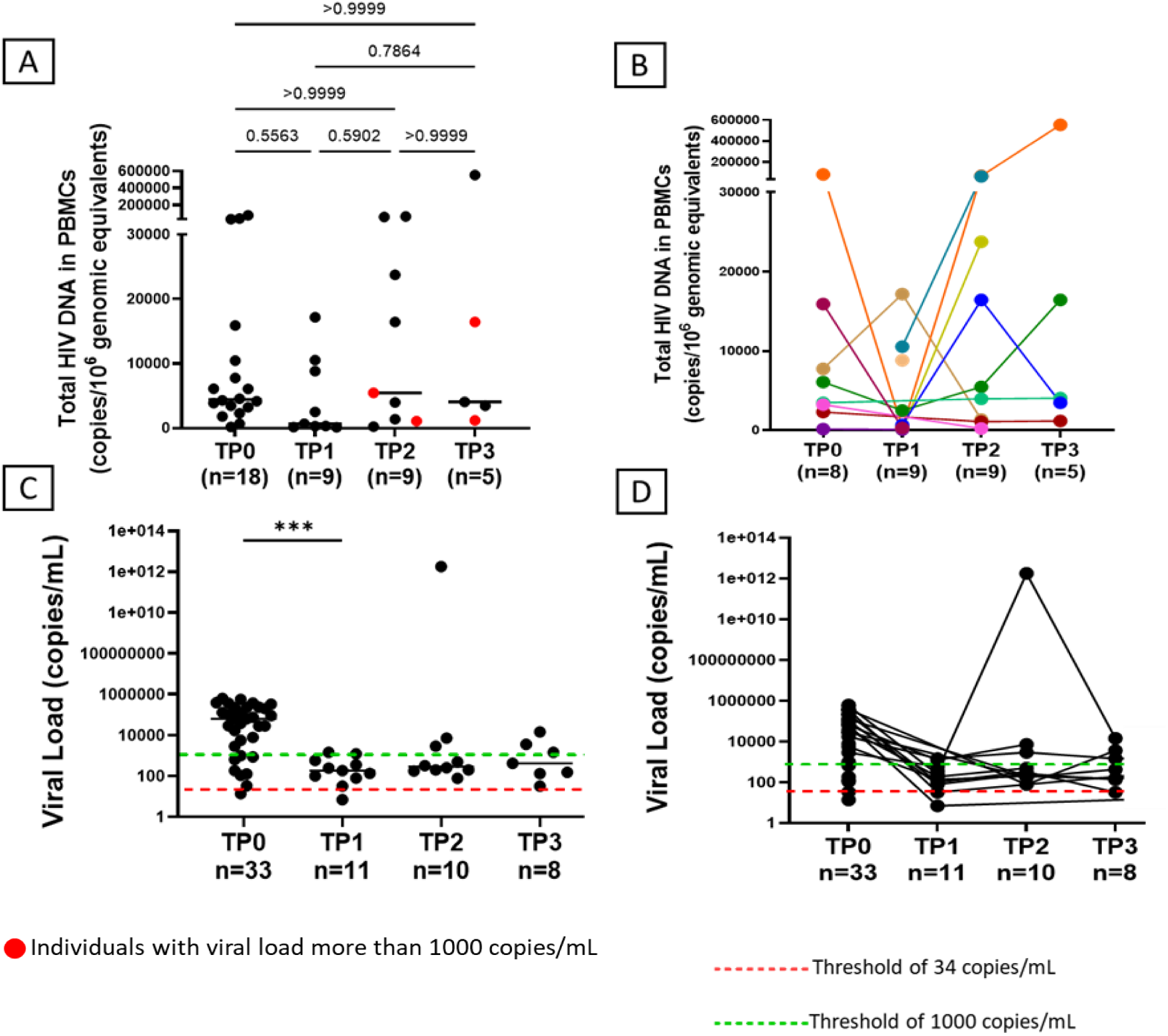
Dynamics of Proviral and viral load within HIV+LTBI+ individuals following ART: A. Unmatched data for proviral loads at TP0, TP1, TP2 and TP3 following initiation of ART. B. Matched data for proviral loads at TP0, TP1, TP2 and TP3 following initiation of ART. C. Unmatched data for viral loads at TP0, TP1, TP2 and TP3 following initiation of ART. D. Matched data for viral loads at TP0, TP1, TP2 and TP3 following initiation of ART. Comparisons between groups were calculated by Kruskal-Wallis one-way ANOVA non-parametric test.

### Drug resistance mutation (DRM) analysis of archival virus at different stages of HIV-TB co-infection

HIV quasispecies diversity may be further amplified in individuals co-infected with tuberculosis, as TB-associated immune activation and increased viral replication can accelerate viral evolution and promote diversification of resistant variants. We evaluated both the presence and relative frequency of drug-resistance mutations (DRMs) targeting the four major ART classes i.e. protease inhibitors (PI), nucleoside reverse transcriptase inhibitors (NRTI), non-nucleoside reverse transcriptase inhibitors (NNRTI), and integrase strand transfer inhibitors (INSTI) in archival (cell-associated) proviral DNA, where available, from all three HIV+ groups. The data was obtained from a total of N=39 HIV-1 infected ART naïve individuals further stratified as HIV+LTBI-(n=19), HIV+LTBI+ (n=11) and HIV+TB+ (n=9) individuals. Despite these individuals being ART naïve (as per clinical records) at the time of recruitment we observed a total prevalence (including all drug classes) of 33% DRMs (13/39), occurring at frequencies of 10% or greater across all groups. Of the identified DRMs, 38% (5/13) were attributed to polymorphic mutations associated with INSTIs (Table 2-4). On comparing DRMs between these groups we observed 32% (6/19) DRM within HIV+LTBI-individuals (Figure 4A), 55% (6/11) within HIV+LTBI+ group (Figure 4B) and 11% (1/9) within HIV+TB+ individuals (Figure 4C and Table 4) suggestive of highest DRM occurrence in HIV+LTBI+ individuals compared to the group without co-infection. Surprisingly, PLHIV with active TB infection exhibited low frequency of DRM compared to other groups probably because these individuals had the least circulating CD4+ T cell counts (Table 1) and thus infected cells due to advanced disease progression. In case of HIV+LTBI-individuals, we observed around 10% potential low level of resistance against INSTIs ranging from 31-100% of frequency of mutation and 21% resistance against NNRTIs wherein 2 individuals (IU-0027 and IU-0054) exhibited high level of resistance against Efavirenz (EFV) and Nevirapine (NVP) with 99% of K103N mutation (Table 2). Interestingly, individual IU-0051 and IU-0152 showed low level resistance and high level of resistance respectively against all NNRTIs except Doravirine (DOR) (Table 2).

**Figure 4.**
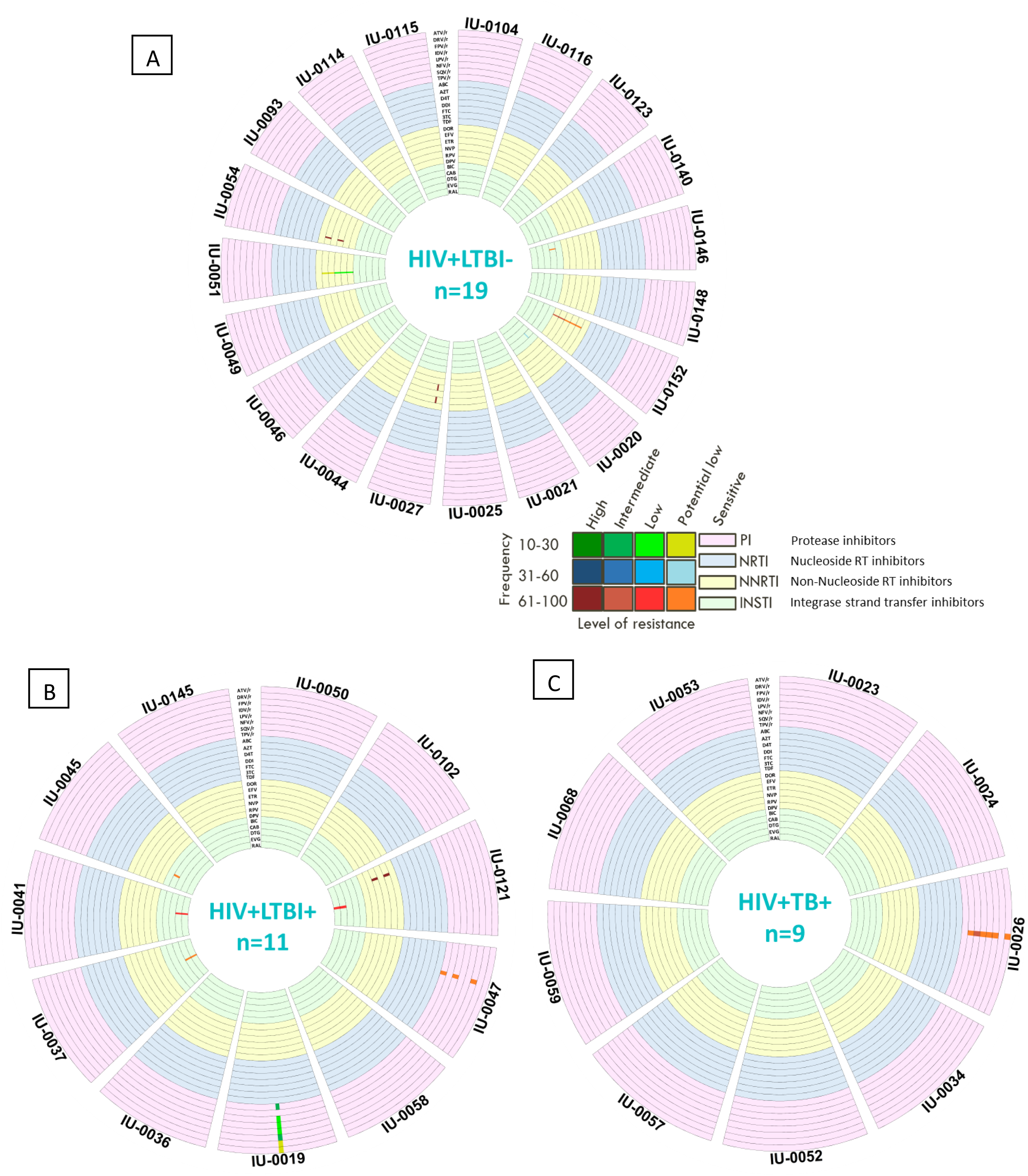
Drug resistance mutation (DRM) analysis of archival viruses: Circos heatmap of residue wise drug resistance mutations (DRMs) in pol sequences in (A) HIV+LTBI-group (n=19), (B) HIV+LTBI+ group (n = 11), and (C) HIV+TB+ group (n = 9). Selection criteria of read depth >50 and mutation detection threshold >10% applied to predict DRMs using the HIVDB tool. Drug resistance summary represented in circos format with a total of 23 concentric tracks of the heatmap representing 23 ARVs from PI (pink), NRTI (blue), NNRTI (yellow), INSTI (green) classes viz from periphery to center (PI-ATV/r, DRV/r, FPV/r, IDV/r, LPV/r, NFV, SQV/r, TPV/r; NRTI-ABC, AZT, D4T, DDI, FTC, 3TC, TDF; NNRTI-DOR, EFV, ETR, NVP, RPV, DPV; INSTI-BIC, CAB, DTG, EVG, and RAL). Each section represents a participant with its ID. Mutations are shown as colored lines indicating their relative position in the gene, level of resistance, and frequency of mutation as described in the key.

**Table 2:** Detailed table for DRM from archival virus in HIV+LTBI-group.

| Sequence Name | Gene | Drug category | Drug name | Resistance Mutation | Level of Resistance | Frequency of mutation | Type of mutation |
| --- | --- | --- | --- | --- | --- | --- | --- |
| IU-0146 | IN | INSTI | CAB | L74M | Potential low | 99.67 | Polymorphic mutation |
| IU-0152 | RT | NNRTI | EFV | E138K | Potential low | 79.13 | Non-Polymorphic mutation selected in a high proportion of persons receiving RPV |
|  | RT | NNRTI | ETR | E138K | Potential low | 79.13 |  |
|  | RT | NNRTI | NVP | E138K | Potential low | 79.13 |  |
|  | RT | NNRTI | RPV | E138K | Intermediate | 79.13 |  |
|  | RT | NNRTI | DPV | E138K | Intermediate | 79.13 |  |
| IU-0020 | IN | INSTI | CAB | L74I | Potential low | 60.91 | Polymorphic mutation |
|  | IN | INSTI | CAB | L74M | Potential low | 35.84 | Polymorphic mutation |
| IU-0027 | RT | NNRTI | EFV | K103N | High | 99.10 | Non-polymorphic mutation |
|  | RT | NNRTI | NVP | K103N | High | 99.10 |  |
| IU-0051 | RT | NNRTI | EFV | H221Y | Potential low | 15.83 | Non-polymorphic accessory mutation |
|  | RT | NNRTI | ETR | H221Y | Potential low | 15.83 |  |
|  | RT | NNRTI | NVP | H221Y | Low | 15.83 |  |
|  | RT | NNRTI | RPV | H221Y | Low | 15.83 |  |
|  | RT | NNRTI | DPV | H221Y | Low | 15.83 |  |
| IU-0054 | RT | NNRTI | EFV | K103N | High | 99.57 | Non-polymorphic mutation |
|  | RT | NNRTI | NVP | K103N | High | 99.57 |  |

**Table 3:** Detailed table for DRM in archival virus in HIV+LTBI+ group.

| Sequence Name | Gene | Drug category | Drug name | Resistance Mutation | Level of Resistance | Frequency of mutation | Type of mutation |
| --- | --- | --- | --- | --- | --- | --- | --- |
| IU-0121 | RT | NNRTI | EFV | K103N | High | 96.86 | Non-polymorphic mutation |
|  | RT | NNRTI | NVP | K103N | High | 96.86 |  |
|  | IN | INSTI | EVG | E157Q | Low | 98.91 | Highly polymorphic mutation |
|  | IN | INSTI | RAL | E157Q | Low | 98.91 |  |
|  | IN | INSTI | RAL | G163R | Low | 88.15 | Non-polymorphic mutation |
|  | IN | INSTI | EVG | G163R | Low | 88.15 |  |
| IU-0047 | PR | PI | FPV/r | L33F | Potential low | 99.23 | Non-polymorphic accessory mutation |
|  | PR | PI | NFV | L33F | Potential low | 99.23 |  |
|  | PR | PI | TPV/r | L33F | Potential low | 99.23 |  |
| IU-0019 | PR | PI | ATV/r | I47V | Potential low | 13.56 | Non-polymorphic PI-selected mutation |
|  | PR | PI | DRV/r | I47V | Potential low | 13.56 |  |
|  | PR | PI | FPV/r | I47V | Intermediate | 13.56 |  |
|  | PR | PI | IDV/r | I47V | Low | 13.56 |  |
|  | PR | PI | LPV/r | I47V | Low | 13.56 |  |
|  | PR | PI | NFV | I47V | Low | 13.56 |  |
|  | PR | PI | TPV/r | I47V | Intermediate | 13.56 |  |
| IU-0037 | IN | INSTI | EVG | E157Q | Potential low | 99.13 | Highly polymorphic mutation |
|  | IN | INSTI | RAL | E157Q | Potential low | 99.13 |  |
| IU-0041 | IN | INSTI | EVG | G163K | Low | 98.65 | Non-polymorphic mutation |
|  | IN | INSTI | RAL | G163K | Low | 98.65 |  |
| IU-0045 | IN | INSTI | CAB | L74M | Potential low | 64.06 | Polymorphic mutation |

**Table 4:**
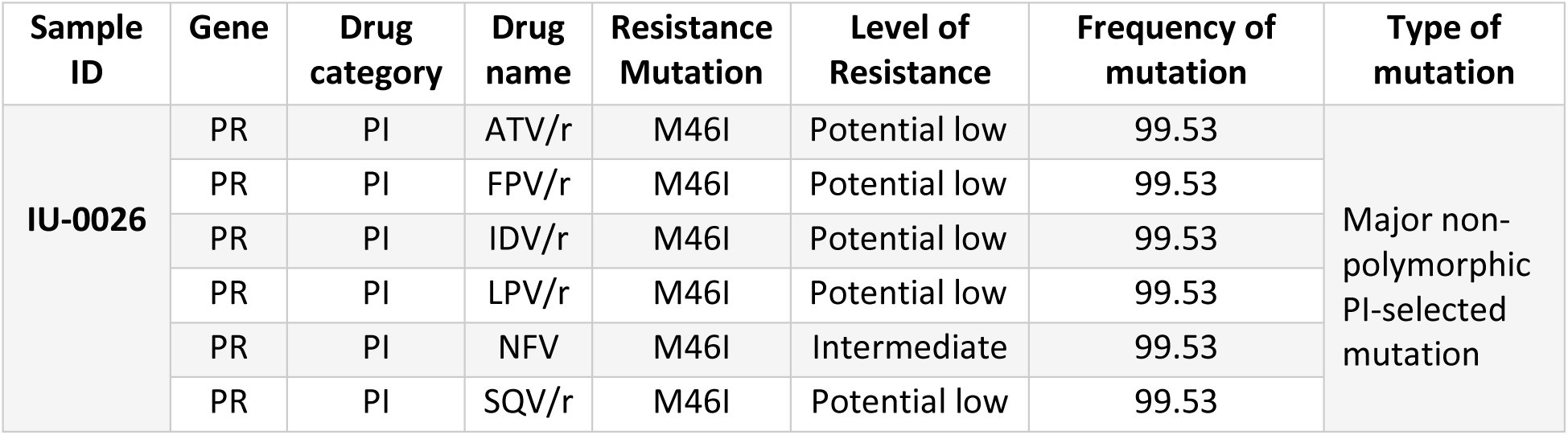
Detailed table for DRM from archival virus in HIV+TB+ group.

| Sample ID | Gene | Drug category | Drug name | Resistance Mutation | Level of Resistance | Frequency of mutation | Type of mutation |
| --- | --- | --- | --- | --- | --- | --- | --- |
| IU-0026 | PR | PI | ATV/r | M46I | Potential low | 99.53 | Major non-polymorphic PI-selected mutation |
|  | PR | PI | FPV/r | M46I | Potential low | 99.53 |  |
|  | PR | PI | IDV/r | M46I | Potential low | 99.53 |  |
|  | PR | PI | LPV/r | M46I | Potential low | 99.53 |  |
|  | PR | PI | NFV | M46I | Intermediate | 99.53 |  |
|  | PR | PI | SQV/r | M46I | Potential low | 99.53 |  |

HIV+LTBI+ individuals exhibited 36% (4/11) resistance against INSTIs, 9% (1/11) resistance against NNRTIs along with 18% (2/11) resistance against PIs. Out of the 4 individuals who showed resistance against INSTIs 3 individuals (IU-0037, IU-0041 and IU-0121) exhibited potential low to low level of resistance against Elvitegravir (EVG) and Raltegravir (RAL) with mutation E157Q at position 157, G163K and G163R at position 163 ranging from 61-100%. Notably, individuals IU-0019 and IU-0047 exhibited resistance against PIs wherein IU-0019 showed potential low to intermediate level resistance due to I47V mutation with a frequency of 13% against all the PIs except for Saquinavir (SQV) (Table 3). IU-0047 displayed potential low level of resistance against Tipranavir (TPV), Nelfinavir (NFV) and Fosamprenavir (FPV) with 99% of mutation of L33F at position 33 (Table 3). It was interesting to note that individuals in HIV+LTBI- and HIV+LTBI+ groups exhibited resistance against the class of drugs (INSTIs) included in the current 1st and 2^nd^ line ART regimen that was never introduced before.

### Drug resistance mutation (DRM) analysis of circulating virus at different stages of HIV-TB co-infection

We next evaluated DRMs in circulating viruses from plasma samples reflective of the actively replicating viral population, which directly influences treatment efficacy. DRMs were analysed from (n=7) HIV+LTBI-, (n=5) HIV+LTBI+ and (n=9) HIV+TBI+ individuals. We observed a total prevalence (including all drug classes) of 33% (7/21) DRMs which is similar to observation from archival viruses at frequencies of more than 10% across all groups. These DRMs predominantly involved INSTI resistance, accounting for 71% (5/7), and were entirely attributed to polymorphic mutations associated with the INSTI class (Table 5-7) while the remaining 29% (2/7) was against NNRTI class of drugs. The occurrence of group wise DRM distribution observed was, 43% (3/7) within HIV+LTBI-individuals (Figure 5A), 40% (2/5) within HIV+LTBI+ group (Figure 5B) and 22% (2/9) within HIV+TB+ individuals (Figure 5C). The individuals IU-0036, IU-0041 (HIV+LTBI+ group) and IU-0062 (HIV+TB+ group) exhibited potential low to low level of resistance mutations with a mutation frequency of 99% against class of INSTIs (EVG (Elvitegravir) and RAL (Raltegravir) drugs) due to E157Q and G163K mutations (Table 6 and 7). Individuals IU-0131 and IU-0151 from HIV+LTBI-group showed potential low to intermediate level of resistance mutations against all the drugs from NNRTI class except for DOR (Doravirine) with a mutation frequency of 31-100% at E138A and V179D (Table 5).

**Figure 5.**
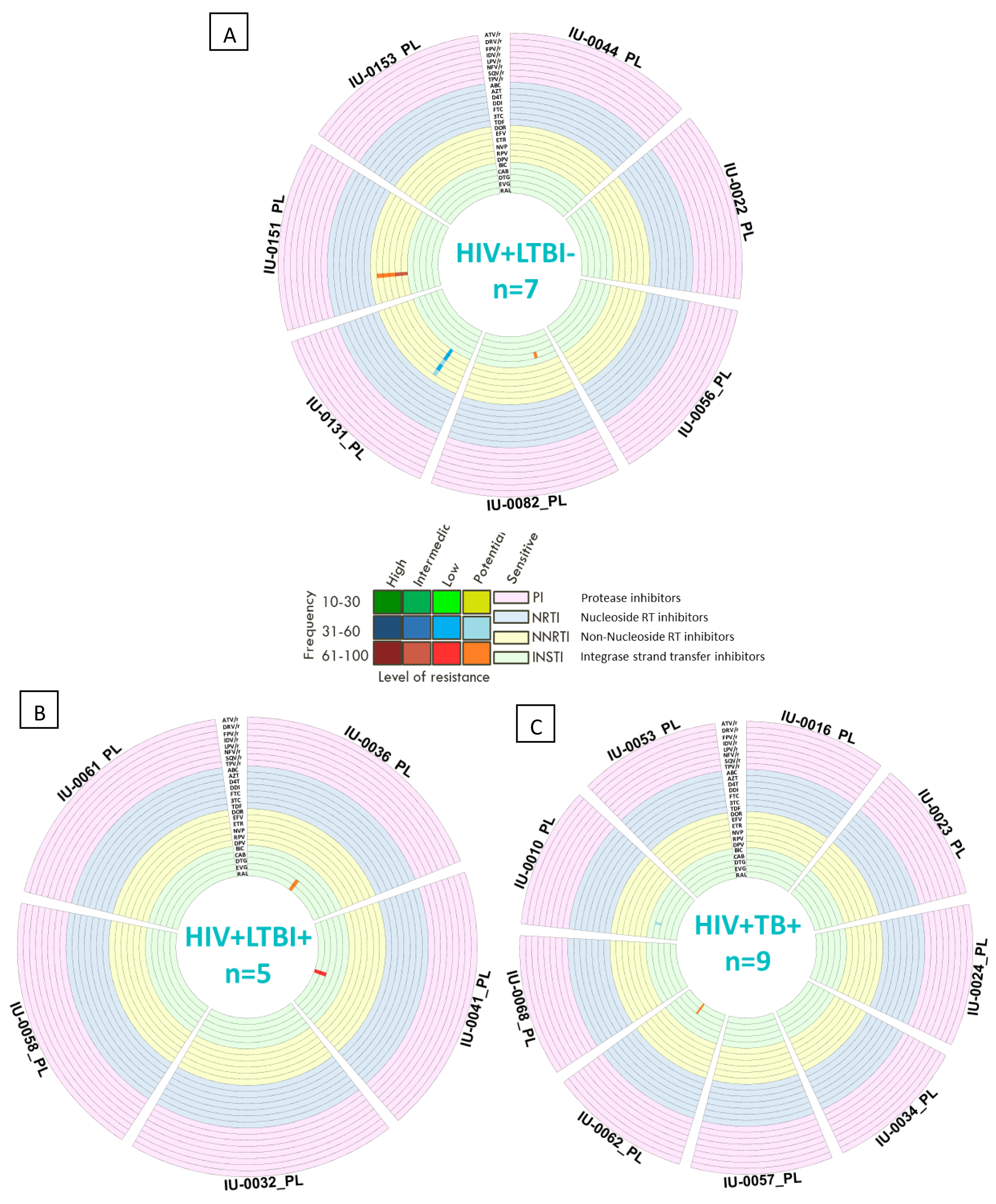
Drug resistance mutation (DRM) analysis of circulating viruses: Circos heatmap of residue wise drug resistance mutations (DRMs) in pol sequences in (A) HIV+LTBI-group (n=8), (B) HIV+LTBI+ group (n = 5), and (C) HIV+TB+ group (n = 8). Selection criteria of read depth >50 and mutation detection threshold >10% applied to predict DRMs using the HIVDB tool. Drug resistance summary represented in circos format with a total of 23 concentric tracks of the heatmap representing 23 ARVs from PI (pink), NRTI (blue), NNRTI (yellow), INSTI (green) classes viz from periphery to center (PI-ATV/r, DRV/r, FPV/r, IDV/r, LPV/r, NFV, SQV/r, TPV/r; NRTI-ABC, AZT, D4T, DDI, FTC, 3TC, TDF; NNRTI-DOR, EFV, ETR, NVP, RPV, DPV; INSTI-BIC, CAB, DTG, EVG, and RAL). Each section represents a participant with its ID. Mutations are shown as colored lines indicating their relative position in the gene, level of resistance, and frequency of mutation as described in the key.

**Table 5:** Detailed table for DRM from circulating virus in HIV+LTBI-group.

| Sample ID | Gene | Drug category | Drug name | Resistance mutation | Level of mutation | Frequency of mutation | Type of mutation |
| --- | --- | --- | --- | --- | --- | --- | --- |
| IU-0131_PL | RT | NNRTI | EFV | E138A | Potential low | 31.63 | Non-polymorphic mutation |
|  | RT | NNRTI | ETR | E138A | Low | 31.63 |  |
|  | RT | NNRTI | NVP | E138A | Potential low | 31.63 |  |
|  | RT | NNRTI | RPV | E138A | Low | 31.63 |  |
|  | RT | NNRTI | DPV | E138A | Low | 31.63 |  |
| IU-0151_PL | RT | NNRTI | EFV | E138K | Potential low | 99.66 | Non-Polymorphic mutation selected in a high proportion of persons receiving RPV |
|  | RT | NNRTI | ETR | E138K | Potential low | 99.66 |  |
|  | RT | NNRTI | NVP | E138K | Potential low | 99.66 |  |
|  | RT | NNRTI | RPV | E138K | Intermediate | 99.66 |  |
|  | RT | NNRTI | DPV | E138K | Intermediate | 99.66 |  |
| IU-0082_PL | IN | INSTI | CAB | L74I | Potential low | 98.97 | Highly polymorphic mutation |

**Table 6:**
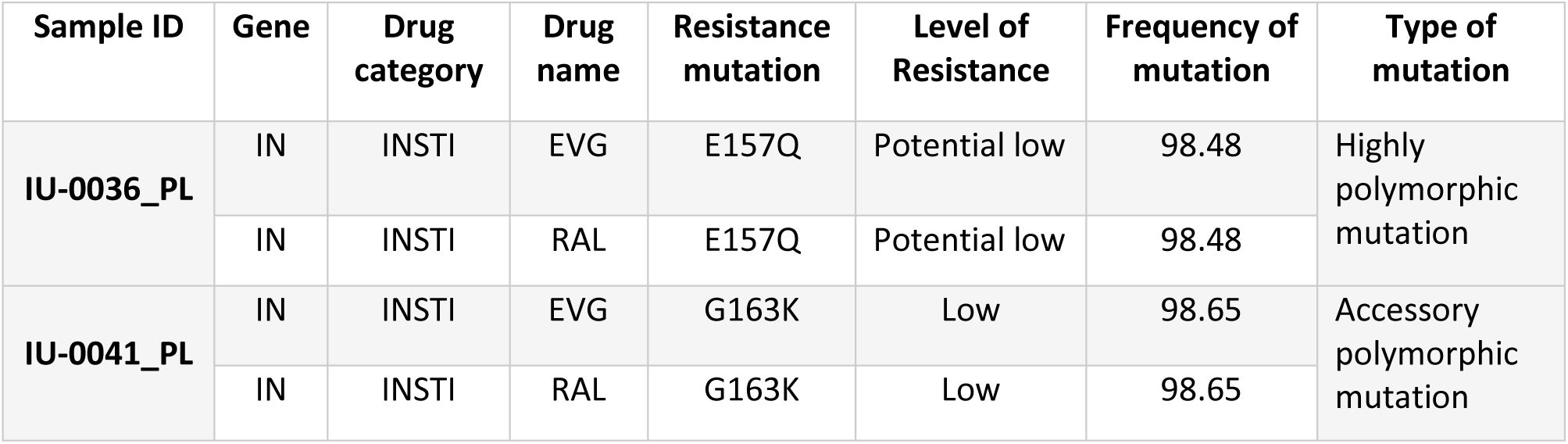
Detailed table for DRM from circulating virus in HIV+LTBI+ group.

| Sample ID | Gene | Drug category | Drug name | Resistance mutation | Level of Resistance | Frequency of mutation | Type of mutation |
| --- | --- | --- | --- | --- | --- | --- | --- |
| IU-0036_PL | IN | INSTI | EVG | E157Q | Potential low | 98.48 | Highly polymorphic mutation |
|  | IN | INSTI | RAL | E157Q | Potential low | 98.48 |  |
| IU-0041_PL | IN | INSTI | EVG | G163K | Low | 98.65 | Accessory polymorphic mutation |
|  | IN | INSTI | RAL | G163K | Low | 98.65 |  |

**Table 7:**
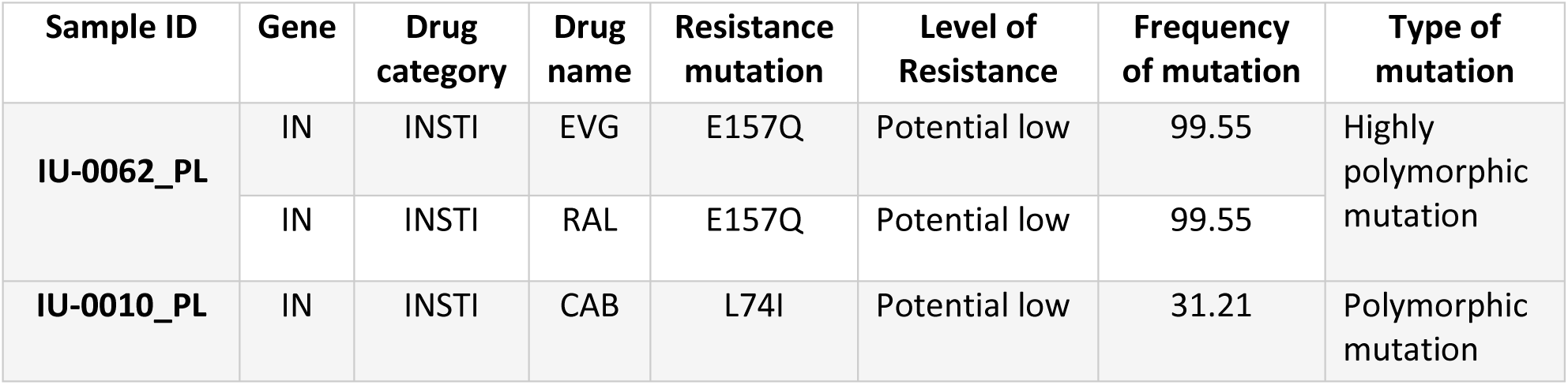
Detailed table for DRM from circulating virus in HIV+TB+ group.

Analysis of matched data from plasma (Figure 6 A) and PBMCs (Figure 6 B) from 10 individuals across all the groups, showed concordance for all the samples except for IU-0036 which exhibited potential low-level DRM against INSTIs class of drugs suggesting that the mutation existed in the currently replicating viral population but was not yet archived in the cellular reservoir or present at very low (undetectable) levels.

**Figure 6.**
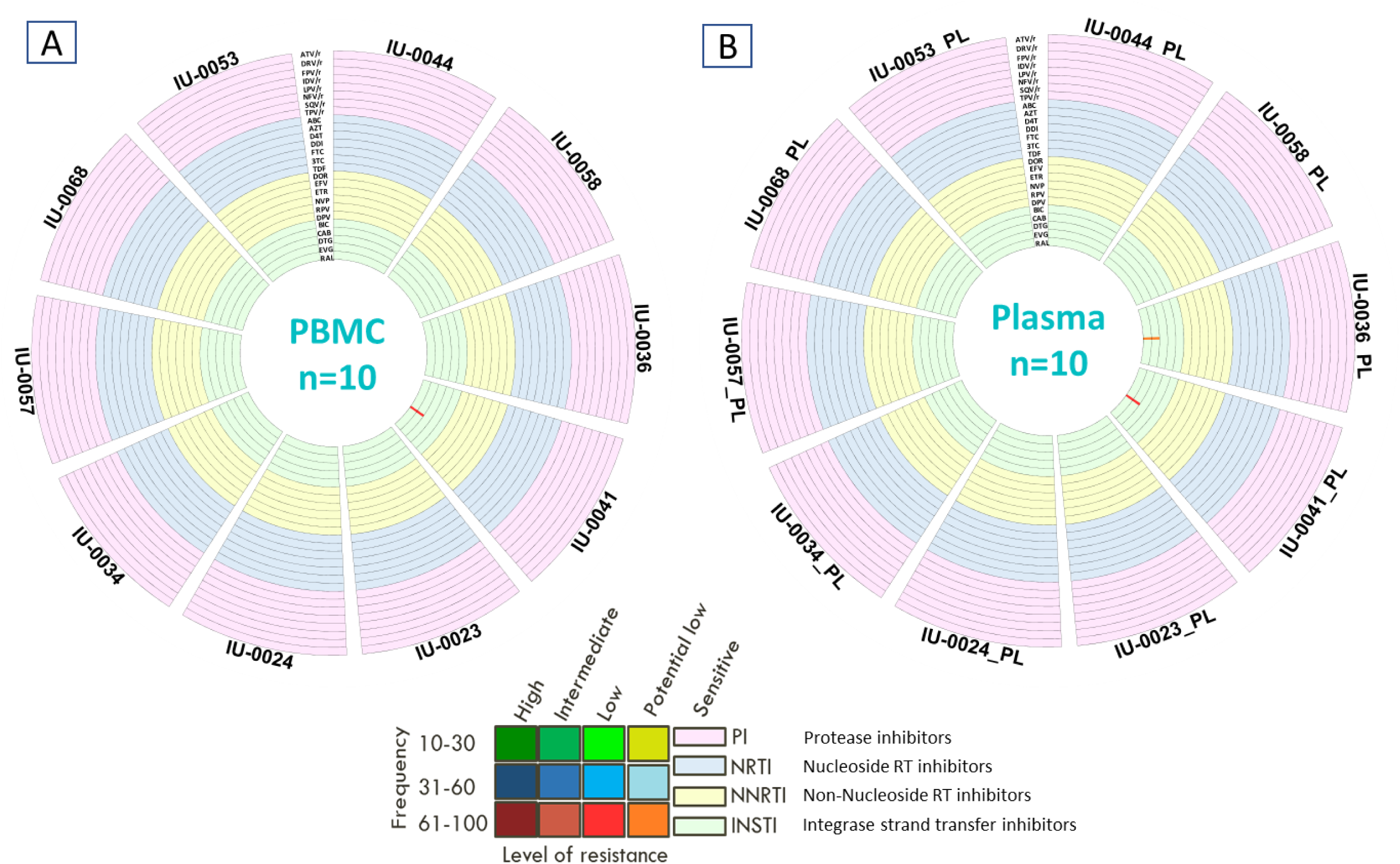
Comparison of Drug resistance mutation (DRM) analysis in matched archival and circulating viruses: Circos heatmap of residue wise drug resistance mutations (DRMs) in pol sequences in (A) PBMCs (n=10) and (B) Plasma group (n = 10). Selection criteria of read depth >50 and mutation detection threshold >10% applied to predict DRMs using the HIVDB tool. Drug resistance summary represented in circos format with a total of 23 concentric tracks of the heatmap representing 23 ARVs from PI (pink), NRTI (blue), NNRTI (yellow), INSTI (green) classes viz from periphery to center (PI-ATV/r, DRV/r, FPV/r, IDV/r, LPV/r, NFV, SQV/r, TPV/r; NRTI-ABC, AZT, D4T, DDI, FTC, 3TC, TDF; NNRTI-DOR, EFV, ETR, NVP, RPV, DPV; INSTI-BIC, CAB, DTG, EVG, and RAL). Each section represents a participant with its ID. Mutations are shown as colored lines indicating their relative position in the gene, level of resistance, and frequency of mutation as described in the key.

### Phylogenetic analysis of env diversity at different stages of HIV-TB coinfection

Phylogenetic analysis of HIV-1 env sequences was performed to assess viral genetic diversity, infer evolutionary relationships, and examine clustering patterns across HIV+LTBI-, HIV+LTBI+, and HIV+TB+ groups (Figure 7). No distinct clustering was observed among the three HIV+ groups, and analysis of average pairwise distance (APD) revealed no significant differences between HIV+LTBI-, HIV+LTBI+ and HIV+TB+ individuals (Figure 8A). This suggests largely overlapping viral populations, particularly between the LTBI- and LTBI+ groups. The apparent lack of samples available for analysis from the HIV+TB+ group probably reflected the difficulty in obtaining env amplicons from these individuals who had very low circulating CD4 counts associated with advanced disease progression. As expected, significantly higher APD values were observed in the PBMC compartment compared to plasma (Figure 8B), suggesting greater genetic diversity within the archival viral reservoir relative to circulating viruses. This observation was further supported by the limited set of matched PBMC and plasma samples (n = 4) (Figure 8C and D). Among these, three pairs demonstrated either monophyletic clustering or close phylogenetic proximity (highlighted in blue rectangles), indicating intra-individual evolutionary relatedness.

**Figure 7.**
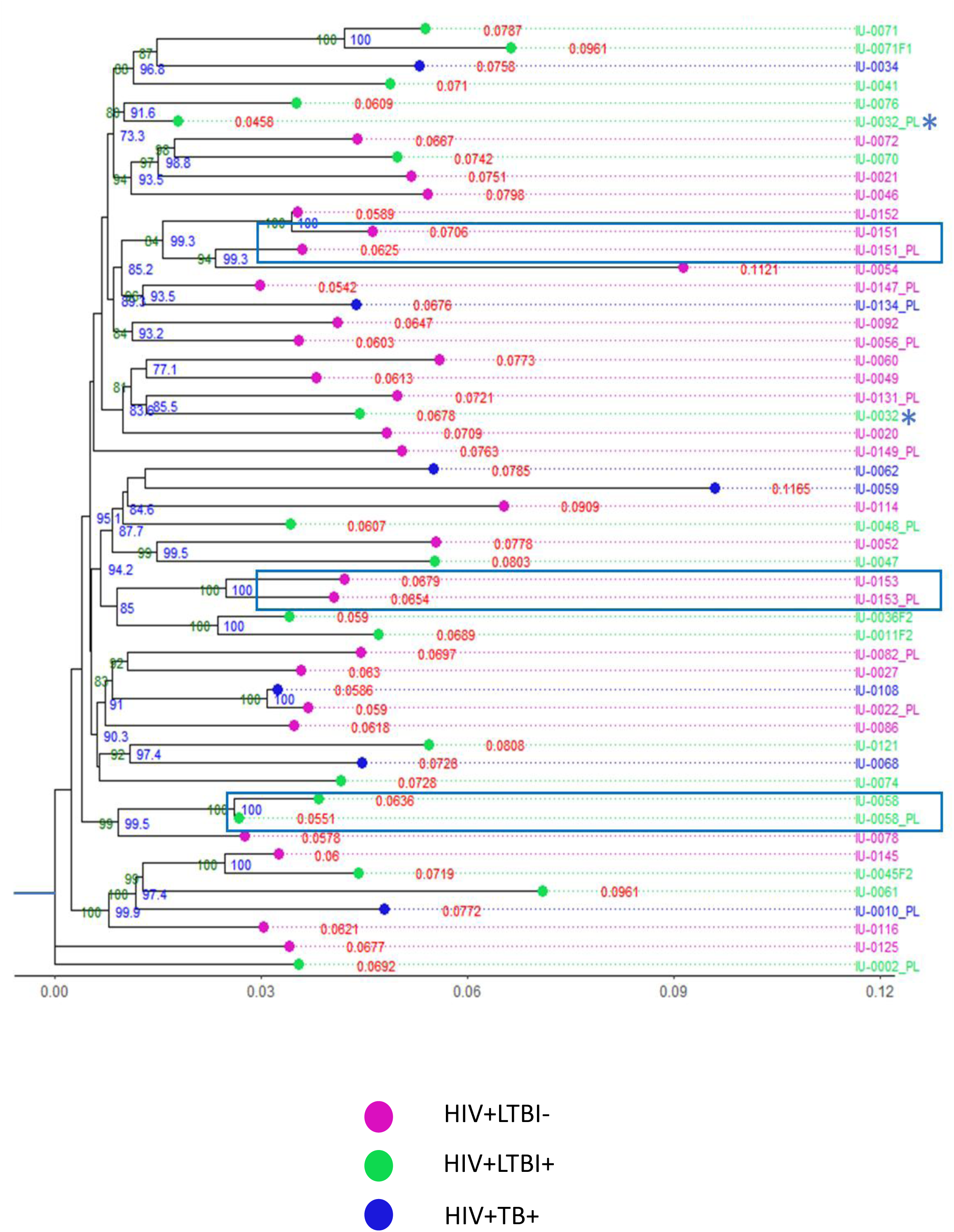
Phylogenetic tree of *env* diversity analysis in different stages of HIV-TB coinfection: Phylogenetic tree of *env* sequences from 3 HIV+ groups. The tree shows clustering on the basis of *env* diversity in sequences obtained from PBMC compartment of 3 groups [HIV+LTBI-(n=19), HIV+LTBI+ (n=10) and HIV+TB+ (n=5)], Plasma compartment (denoted as PL next to sequence ID; n=14 across all 3 groups) and follow-up sequences (denoted as F1/F2 next to sequence ID; n=4). The sequence IDs highlighted in blue represents matched PBMC and plasma sequences showing monophyletic or close clustering. Values in red represent average pairwise distance (APD), values in blue represent bootstrap values and values in green represent SH-aLRT values. gg tree package in R studio was used to generate the phylogenetic tree. Sequence marked with * indicate matched PBMC and plasma data that did not show close phylogenetic clustering.

**Figure 8.**
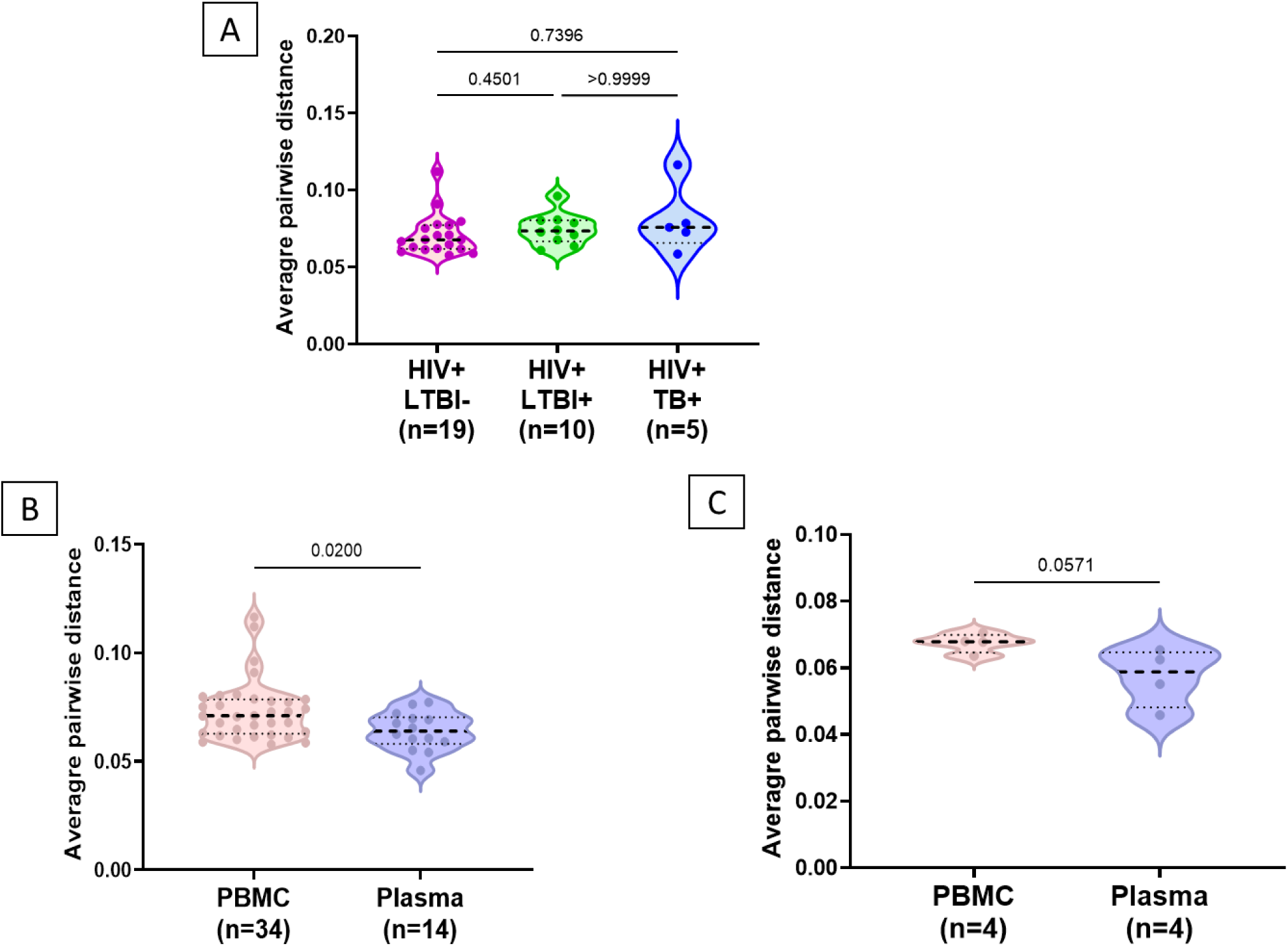
Analysis of Average Pairwise Distance (APD) from phylogenetic tree: Difference in APD values (A) between *env* sequences in HIV+LTBI-, HIV+LTBI+ and HIV+TB+ groups (B) between *env* sequences in PBMC and plasma compartment (C) between *env* sequences in matched PBMC and plasma compartment. Comparisons between groups were performed by Kruskal–Wallis one-way ANOVA non-parametric test and comparison between PBMC and plasma compartment for matched samples was done using Mann-Whitney t-test for unpaired analysis and Wilcoxon signed rank test for paired analysis.

## Discussion

The persistence of Human immunodeficiency virus (HIV) despite effective antiretroviral therapy (ART) remains one of the major barriers to achieving a functional cure (21). The persistence of long-lived latently infected cells collectively referred to as the HIV reservoir enables viral rebound upon treatment interruption and therefore represents the principal obstacle to HIV eradication strategies (21–23). Among PLHIV, TB remains the most common opportunistic infection and a major cause of morbidity and mortality globally (24). Studies have reported that episodes of tuberculosis can lead to an increase in plasma HIV viral load, even among individuals who are undergoing antiretroviral therapy (2,6). Hence, understanding how TB co-infection, particularly LTBI, influences HIV reservoir size and dynamics is therefore critical for designing effective HIV cure strategies.

In the present study, we examined proviral DNA levels, representative of the circulating HIV-1 reservoir, in ART-naïve HIV-1C infected individuals across three clinical groups: HIV+LTBI−, HIV+LTBI+, and HIV+TB+. Proviral DNA serves as a widely accepted surrogate marker for the latent HIV reservoir, as it reflects integrated viral genomes within host cells that can potentially produce replication-competent virus (3,25). Our results indicated that although proviral loads did not differ significantly across groups, a trend toward higher proviral loads in PLHIV co-infected with active TB was observed. This observation may be plausible as active TB infection induces strong systemic immune activation characterized by increased production of inflammatory cytokines such as TNF-α, IL-6 and IFN-γ (11,26). Also, chronic immune activation is known to enhance HIV replication and promote infection of activated CD4+ T cells, thereby potentially expanding the size of the viral reservoir (27–30).

We also observed an interesting correlation between systemic immune markers with the reservoir size in HIV+LTBI+ individuals compared to PLHIV with active TB co-infection or with the ones without any co-infection suggestive of the role of LTBI in modulating HIV persistence as latent TB infection represents a chronic immunological stimulus even in the absence of active disease (31–33). Studies have shown that individuals with LTBI exhibit persistent antigen-specific immune responses and low-grade immune activation due to the continued presence of mycobacterial antigens (28,33,34). Such immune activation may influence HIV reservoir dynamics by promoting periodic activation of latently infected cells, potentially contributing to low-level viral transcription or clonal expansion of infected CD4+ T cells. Therefore, investigating HIV reservoir size in individuals with LTBI is particularly relevant in TB-endemic settings, where a large fraction of PLHIV may harbor latent TB infection.

The study also provides insights into the landscape of drug resistance mutations (DRMs) and viral evolution in both proviral DNA and circulating plasma virus across different stages of HIV–TB co-infection in ART-naïve individuals. Despite the absence of prior antiretroviral exposure, an overall DRM prevalence of 33% was observed in both compartments, with 38% and 71% polymorphic mutation associated with INSTI in PBMC and plasma derived viruses respectively. This supports the concept that certain INSTI-associated mutations, such as E157Q, G163K/R, and L74I, can arise as naturally occurring polymorphisms (35,36). These polymorphic mutations are often present in ART-naïve populations and typically confer minimal or no resistance on their own, but they may facilitate the emergence or enhance the impact of major resistance mutations under drug pressure (37) of other drugs from same class such as Dolutegravir (DTG) which is a part of current 1st line therapy regimen. In contrast, the detection of non-polymorphic mutations, such as K103N and E138A, against NNRTIs is of greater clinical concern as these mutations are strongly associated with transmitted drug resistance (TDR). For example, K103N, observed at high frequency (99%) in our cohort, is a well-established major NNRTI resistance mutation conferring high-level resistance to Efavirenz (EFV) and Nevirapine (NVP). The presence of such mutations in ART-naïve individuals indicates transmission of resistant viral strains, which can significantly compromise treatment efficacy if regimens containing these drugs (either standard or alternate) are initiated without prior resistance testing (38) and continuous surveillance (3,20). Our observation of DRM analysis aligns with previous reports indicating that primary resistance mutations can arise in untreated individuals and may compromise therapy outcomes (39,40).

As observed in our proviral DNA analysis, we found an underrepresentation of sequences obtained from our actively coinfected group. This could have been due to reduced CD4+ T cell availability (12), and consequently fewer analysable infected cells. While other studies claim to have observed greater DRM in HIV-TB coinfected individuals compared to non-coinfected group (41), this difference could be attributed to better median CD4+ T cell counts in HIV+TB+ group compared to our cohort. A notable finding was the higher prevalence of DRMs in HIV+LTBI+ individuals compared to HIV+LTBI-group. This may reflect a state of underlying chronic immune activation without the severe CD4 depletion (12,42), allowing sustained viral replication and diversification. In addition to systemic immune activation, the higher prevalence of DRMs observed in HIV+LTBI+ individuals may also be attributed to the presence of tuberculous granulomas, which act as distinct anatomical and immunological niches for viral persistence and replication (17,43). In the context of HIV infection, these sites can harbour infected CD4+ T cells and macrophages, thereby providing a localized reservoir for ongoing viral replication and evolution (42,44) and our results show for the first time in primary samples, derived from infected individuals, that this may reflect such physiological processes. Importantly, the immune milieu within granulomas differs significantly with restricted immune surveillance, and variable ATT drug penetration (44,45). Such conditions may promote independent viral evolution and selection of variants, thereby contributing to the emergence of DRMs even in the absence of ART.

Our findings also demonstrated a high concordance of 90% between the two compartments in a limited cohort, which is consistent with previous reports indicating a median concordance rate of approximately 87% (38,46–48), including our own study (49), supporting the reliability of PBMC-derived resistance profiles as a proxy for circulating virus. The exception of one individual (IU-0036), where a mutation was detected only in plasma, indicated that the circulating variant may have been derived from another tissue reservoir not represented in the circulating PBMC repertoire. Such compartmental discordance has been previously reported and underscores the importance of analysing both compartments, at least in ART naïve setting for a comprehensive understanding of resistance (50).

In our cohort (n=34) phylogenetic analysis of env sequences did not demonstrate enhanced or distinct clustering among HIV-TB co-infected individuals compared to other groups. In contrast to previous studies (17) reporting increased diversification in actively co-infected populations. This lack of observable difference may be due the difficulty in obtaining env amplicons from these individuals who had very low circulating CD4 counts due to advanced disease progression. Overall, the significantly higher diversity observed in PBMC-derived sequences is consistent with previous reports demonstrating that reservoirs harbour genetically diverse archival variants accumulated over time (51,52).

In summation, these results highlight the possible presence of substantial transmitted drug resistance and concordance of resistance profiles across plasma and PBMC compartments in ART-naïve individuals. Also, DRM results from LTBI infected cohort suggests that there indeed may be contribution of increased viral diversification due to coinfection with Latent TB. Additionally, the enhanced diversity within archival reservoirs underscores their role in shaping HIV evolution. These findings emphasize the importance of baseline resistance screening and comprehensive compartmental analysis together with continued surveillance to achieve effective treatment strategies.

## Author Contributions

Conceptualization: V.P. (Vainav Patel), N.M. (Nupur Mukherjee), K.M., T.M. and V.M.B.; methodology: S.B. (Shilpa Bhowmick), S.Y., K.K., P.K., S.S., P.D., S.K. (Snehal Kaginkar), V.P. (Varsha Padwal), N.N., S.M., J.S. (Jyoti Sutar) and N.M. (Nandan Mohite); formal analysis: S.B. (Shilpa Bhowmick), S.B. (Sharad Bhagat) and J.S. (Jyoti Sutar); investigation: S.B. (Shilpa Bhowmick); Facilitation of participant recruitment: V.N., P.P., S.A., S.G. and J.S. (Jayanthi Shastri); resources: V.P. (Vainav Patel); data curation: S.B. (Shilpa Bhowmick)and S.B. (Sharad Bhagat); writing—original draft preparation: S.B. (Shilpa Bhowmick) and V.P. (Vainav Patel); writing—review and editing, S.B. (Shilpa Bhowmick); V.P. (Vainav Patel) and J.B.; supervision: V.P. (Vainav Patel); project administration: T.M., S.B. (Shilpa Bhowmick), P.D., J.S. (Jyoti Sutar), J.B. and V.P. (Vainav Patel); funding acquisition: N.M. (Nupur Mukherjee), K.M., V.P. (Vainav Patel), T.M. and J.B. All authors have read and agreed to the published version of the manuscript.

## Acknowledgments

We are grateful to the study participants and to the National AIDS control Organisation (NACO), Ministry of Health & Family Welfare, Govt. of India for facilitating recruitment in this study. We appreciate help and support from Chromosome Laboratory Pvt. Ltd. for their sequencing services.

